# Exogenous putrescine alleviates photoinhibition caused by salt stress through increasing cyclic electron flow in cucumber

**DOI:** 10.1101/301762

**Authors:** Xinyi Wu, Sheng Shu, Yu Wang, Ruonan Yuan, Shirong Guo

## Abstract

When plants suffer from abiotic stresses, cyclic electron flow (CEF) is induced for photoprotection. Putrescine (Put), a main polyamine in chloroplasts, plays a critical role in stress tolerance. To elucidate the mechanism of Put regulating CEF for salt-tolerance in cucumber leaves, we measured chlorophyll fluorescence, P700 redox state, ATP and NADPH accumulation and so on. The maximum photochemical efficiency of PSII (Fv/Fm) was not influenced by NaCl and/or Put, but the activity of PSI reaction center (P700) was seriously inhibited by NaCl. Salt stress induced high level of CEF, moreover, NaCl and Put treated plants exhibited much higher CEF activity and ATP accumulation than single salt-treated plants to provide adequate ATP/NADPH ratio for plants growth. Furthermore, Put decreased the trans-membrane proton gradient (ΔpH), accompanied by reducing the pH-dependent non-photochemical quenching (qE) and increasing efficient quantum yield of PSII (Y(II)). The ratio of NADP^+^/NADPH in salt stressed leaves was significantly increased by Put, indicating that Put relieved over-reduction pressure at PSI accepter side. Taken together, our results suggest that exogenous Put enhances CEF to supply extra ATP for PSI recovery and CO_2_ assimilation, decreases ΔpH for electron transport related proteins staying active, and enable the non-photochemical quenching transformed into photochemical quenching.

## Introduction

In recent years, secondary salinization has become a major environmental factor that limits yield and quality of vegetable crops in protected cultivation of China. Salt stress can induce ionic toxicity and osmotic stress, which result in the damage to biological macromolecules and interfer with metabolisms in plant cells (Munns and Tester, 2008; Yan et al., 2015). Photosynthesis fuels plant growth, but it is sensitive to salt stress, by virtue of this correlation, photosynthetic capacity is thought to be an important criterion for diagnosing plant adaptability to salinity (Kalaji et al., 2011).

In the process of plants evolution, the photosynthetic thylakoid membrane system has evolved several regulation mechanisms enabling them to adapt to stress condition (Takabayashi et al., 2009). Cyclic eletron flow (CEF) in PSI, an important photoprotection pathway, can regulate electron transport mode to acclimatize itself to adverse environment (Shikanai et al., 1998; Horváth et al., 2000; Munekage et al., 2004). The cyclic process could be visualized as following description: electrons transferred to the PSI reaction center are contributed to primary electron receptor. Then, they are passed to ferredoxin (Fd), an iron containing protein which acts as an electron carrier. The reduced Fd are able to transport electrons to the second electron carrier plastoquinone (PQ) through two available pathways i.e. the protein gradient regulation 5 (PGR5)/PGR-like 1 (PGRL1)-dependent pathway and the NAD(P)H dehydrogenase (NDH) complex-dependent pathway (Burrows et al., 1998; Munekage et al., 2002; Shikanai, 2007; DalCorso et al., 2008). After that, PQ carries electrons to cytochromes *b6f* (Cyt *b6f*). In the process, proton gradient across thylakoid membrane (ΔpH) comes into development, the main functions of ΔpH are: (1) together with Δψ (trans-membrane electric potential) used for ATP synthesis by the ATP synthase; (2) stimulates photoprotection pathway called “non-photochemical quenching”, which competes with photochemical quenching under excessive light conditions and consume the excited energy, avoiding the production of triplet chlorophylls that react with molecular oxygen (Munekage et al., 2002). Ultimately, electrons are returned to P700 by plastocyanin (PC) to finish the cycle.

Recent research has demonstrated that CEF plays an extremely important and essential role in plants photosynthesis and development (Sun et al., 2017). When linear electron flow (LEF) is affected by fluctuant environment, CEF is an essential alternative pathway to replenish ATP and keeps a high ΔpH which is required for energy-dependent quenching (qE, the fast phase of NPQ) to dissipate surplus excited energy (Niyogi, 1999; Muller et al., 2001). Therefore, switching between LEF and CEF is an adaptation mechanism for unsteady environment to keep an appropriate ATP/NADPH ratio required for plants growth. Previous work suggested that LEF plays a dominant role and the effect of CEF is negligible on normal condition (Shikanai, 2007). However, when plants suffer from abiotic stress, LEF is significantly inhibited, and the excess electron preferably injectes to CEF, respiratory chain and combines with O_2_ to produce ROS. Taken together, CEF is indispensable for plants to resist photodamage under stress condition.

Putrescine (Put) is a ubiquitous diamine [NH_2_(CH_2_)_4_NH_2_], which is the obligate precursor of spermidine (Spd) and spermine (Spm). They are low-molecular weight aliphatic amines and constitute the major polyamines in plants (Ioannidis and Kotzabasis, 2007), which are essential for plant growth and development and stress response (Capell et al., 2004; Hussain et al., 2011). Polyamines are firstly recognized as their cationic character, which is one of the main properties believed to mediate their biological activity, therefore they can interact with negatively charged macromolecules such as DNA, RNA, proteins and phospholipids, leading to the stabilization of macromolecules (Kaur-Sawhney et al., 1978; Schuber, 1989; Galstoon and Sawhney, 1990). Some genes involved in PAs biosynthesis are found in the chloroplast, showing their possible relation with photosynthetic process. In chloroplast, PAs conjugating to negatively charged macromolecules is catalyzed by an enzyme named transglutaminase (TGase). This enzyme catalyses the incorporation of PAs into thylakoid and stromal proteins such as the light harvesting complex (LHC) and the large subunit of Rubisco (Hamdani et al., 2011). Besides their polycationic nature, PAs was found playing an other important bioenergetic role as a permeant natural buffer, because they can be found in a charged or uncharged form, although uncharged forms represent less than 0.1%, the physiological role could be crucial in chemiosmosis (Ioannidis and Kotzabasis, 2014). Our previous work has demonstrated that exogenous Put can alleviate the damaging effects of salt stress on the structure and function of the photosynthetic apparatus in salt-stressed cucumber seedling leaves (Shu et al., 2012; Yuan et al., 2014; Yuan et al., 2017). However, the regulation and mechanism of CEF induced by Put has not been reported yet. In the present study, we found Put was more prominent in chemiosmosis regulation, it directly facilitate the production of ATP, which is crucial for P700 repair, the larger content of active P700 induces the higher CEF level; On the other hand, exogenous put is a neutralizer to alleviate reduction of pH caused by CEF. Additinaly appropriate pH in lumen is required for the electron transport related protein to stay active and enable the non-photochemical quenching transformed into photochemical quenching. To sum up, this results reveal an important mechanism for putrescine inducing CEF to alleviate photoinhibition caused by salt stress in cucumber plants, and provides new insights into the role of put in photoprotection to salt stress.

## Materials and methods

### Plant material and growth conditions

Cucumber (*Cucumis sativus* L. Cv. Jinyou NO. 4, obtained from Tian Jin Kernel Cucumber Research Institute, Tianjin, China). seeds were germinated on moist gauze in the dark for 16 h at 28 °C and were placed in 3cm^∗^3cm^∗^4cm sponges. When the first leaves were fully expanded, the seedlings were transplanted into smart climatic chambers (AEtrium 3, AEssense Corp., Shanghai, America) at an irradiance of 400 μmol m^−2^ s^−1^ for 12 h everyday, day/night temperature were set as 28/18°C with ~ 65 ± 5 % relative humidity. The seedlings were grown hydroponically with half-strength Yamazaki soilless culture nutrient solution for cucumber (pH 6.2 ± 0.1, EC 1.0-1.2). When the third leaves were fully expanded, the seedlings were treated as follows: (1) Cont, control, plants were grown in normal condition; (2) Put, normal growth plants were sprayed with 8 mM Put on leaves; (3) NaCl, plants were treated with 90 mM NaCl in nutrient solution; (4) NaCl + Put, salt stressed plants were sprayed with 8 mM NaCl on leaves. Put was sprayed 60 min before the light turned on every day. The third fully expanded leaves (from the top) were sampled after treatment for 7 days and immediately frozen in liquid nitrogen.

### Gas-exchange measurements

Gas-exchange parameters were measured using a portable photosynthesis system (LI-6400, LI-COR Inc, USA) as described by Shu et al (Shu et al., 2013). The photosynthetic rate was measured at 400 ± 10 μmol·mol^−1^ CO^2^, 25°C, 70% relative humidity, and 1500 μmol·m^−2^s^−1^ light intensity.

### Measurement of chlorophyll fluorescence

Chlorophyll (Chl) fluorescence was performed using IMAGING-PAM Chl fluorescence analyser (Heinz Walz, Effeltrich, Germany), plants were fully adapted in dark before measurement. The chlorophyll fluorescense parameters were calculated as follows (Daisuke et al., 2016): maximum photochemical efficiency of PSII, Fv/Fm = (Fm-Fo)Fm; effective photochemical quantum yield of PSII, Y(II) = (Fm’-Fs)/Fm’; non-photochemical quenching, NPQ = *(Fm*-*Fm’)/Fm’;* redox state of Q_A_, 1-qP = 1-(*Fm*’-*F)/(Fm*’-*Fo’*); regulatory energy dissipation quantum yield of PSII, Y(NPQ) = 1-Y(II)-1/(NPQ+1+qL(*Fm*/*Fo*-1)); non-regulatory energy dissipation quantum yield of PSII, Y(NO) = 1/(NPQ+1+qL(*Fm*/*Fo*-1). For photoresponse curve measurement, every intensity light was last 30 seconds. NPQ relaxation was analyzed by following protocol: fully dark adapted plants were illuminated with 396 μmol m^−2^ s^−1^ light for 240 s, followed with 506 s dark, saturated pulse was applied every 20 seconds.

The post-illumination transient increase in Chl fluorescence was determined as previously described (Shikanai et al., 1998). Plants were dark adapted at least 1h, and the optimal room temperature was 25°C, illumination light intensity was set as 396 μmol m^−2^ s^−1^.

### Measurement of P700 redox changes and delayed fluorescence

P700 redox changes was reflected of 820 nm transmission measured by M-PEA (Hansatech, Norfolk, UK) on channel 2. The protocol was modified according to the procedure described by Yan et al (2015). Plants were dark adapted for 30 min, then illuminated with 1s saturated red light (5000 μmol m^−2^ s^−1^), followed with 10s far-red light (intensity = 100%), and the 1 s red light + 10 s far-red light was repeated for 5 times. The P700 oxidation and P700^+^ re-reduction were measured by following procedure: fully dark adapted leaves was firstly induced by far-red light (intensity = 25%, 100s), followed with 30 s dark. Initial rate (0~2s of P700 oxidation; 0~0.3 s of P700^+^ re-reduction) and *t_1/2_* were calculated (Wang et al., 2006).

To calculate ΔI/Io, the first 10 s far-red consequence combined with following formulae were used: Io, the average of the 820-nm transmission signal between 0.4 and 10 ms; Im, the average of the 820-nm transmission signal between 3 and 10 s; ΔI/Io = (Io-Im)/ Io (Zhang et al., 2011).

Delayed fluorescence was measured by M-PEA (Hansatech, Norfolk, UK) on channel 1. Different light intensities are able to present diverse delayed-fluorescence curves, in order to get I_4_, the light intensity was set as 600 μmol m^−2^s^−1^ continued for 200s, (I_1_-D_2_)/D_2_ and (I_4_-D_2_)/D_2_ were calculated as described by Mehta et al. (2011).

### Quantitative real-time PCR

Leaves were light adapted for 20 min before snap frozen in liquid nitrogen, so that the light-induced state can be preserved. RNA was extracted from three biological replicates of cucumber leaves using RNA extraction kit (TIANGEN). cDNA was synthesized from 1 μg of DNase-treated RNA with the TaKaRa First Strand cDNA Synthesis Kit using oligo (dT) primers. Quantitative real-time PCR was performed using gene specific primers (Table S1) in 20 μl reaction system using SYBR Premix Ex Taq II (TaKaRa).

### Determination of light induced ATP, NADPH, NADP^+^ content and activity of RCA

To determine light induced ATP, NADPH and NADP content in leaves, the third leaves which dark adapted at least for 1h and light induced 20 min were collected separately. ATP content was measured as described by He et al. (2015). 0.1 g samples were grinded in ice with 0.9 ml acidity/alkalinity extraction buffer used for NADP^+^/NADPH quantification, the tubes with samples were kept for 5 min at 100 °C in a boiling water. Then, the samples were kept in ice for cooling and centrifugated at 10,000g for 10 min at 4 °C. Supernatant was collected in new tubes and neutralized with the same volume of alkalinity/acidity buffer, centrifugated at 10,000g for 10 min at 4°C, the supernatant was used for analysis. 0.2 g samples were used for tissue homogenate preparation and the RCA activity was measured by ELISA kit.

### Statistical analysis

All biochemical analyses were conducted at least three times. All data were statistically analyzed with SPSS software using Tukey’s test at the P < 0.05 level of significance.

## Results

### Exogenous Put alleviates the inhibition of NaCl on cucumber growth

As shown in Fig. 1A, there was no significant difference between control and Put treated plants under normal conditions. After 7 days of treatment, the growth of NaCl-only treated plants was dramatically inhibited compared with other treatment plants (Fig. 1A). The fresh weight, dry weight and leaf area of NaCl treated plants decreased by 52.24%, 52.78% and 39.96% respectively compared with that in control plants (Fig. S1). In contrast, exogenous Put significantly promoted plant growth in comparison to NaCl-only treated plants (Fig. S1A-C). This trend can also be seen from the data of gas-exchange parameter in Fig. 1B-E that NaCl stress dramatically decreased net photosynthesis rate (Pn), stomata conductance (Gs), intercellular carbon dioxide concentration (Ci) and transpitation rate (Tr) in cucumber leaves, these parameters were recovered by exogenous Put that close to non-stressed leaves. Nevertheless, The results of chlorophyll content showed no obvious different between all treatments (Fig. S1D).

**Fig. 1.**
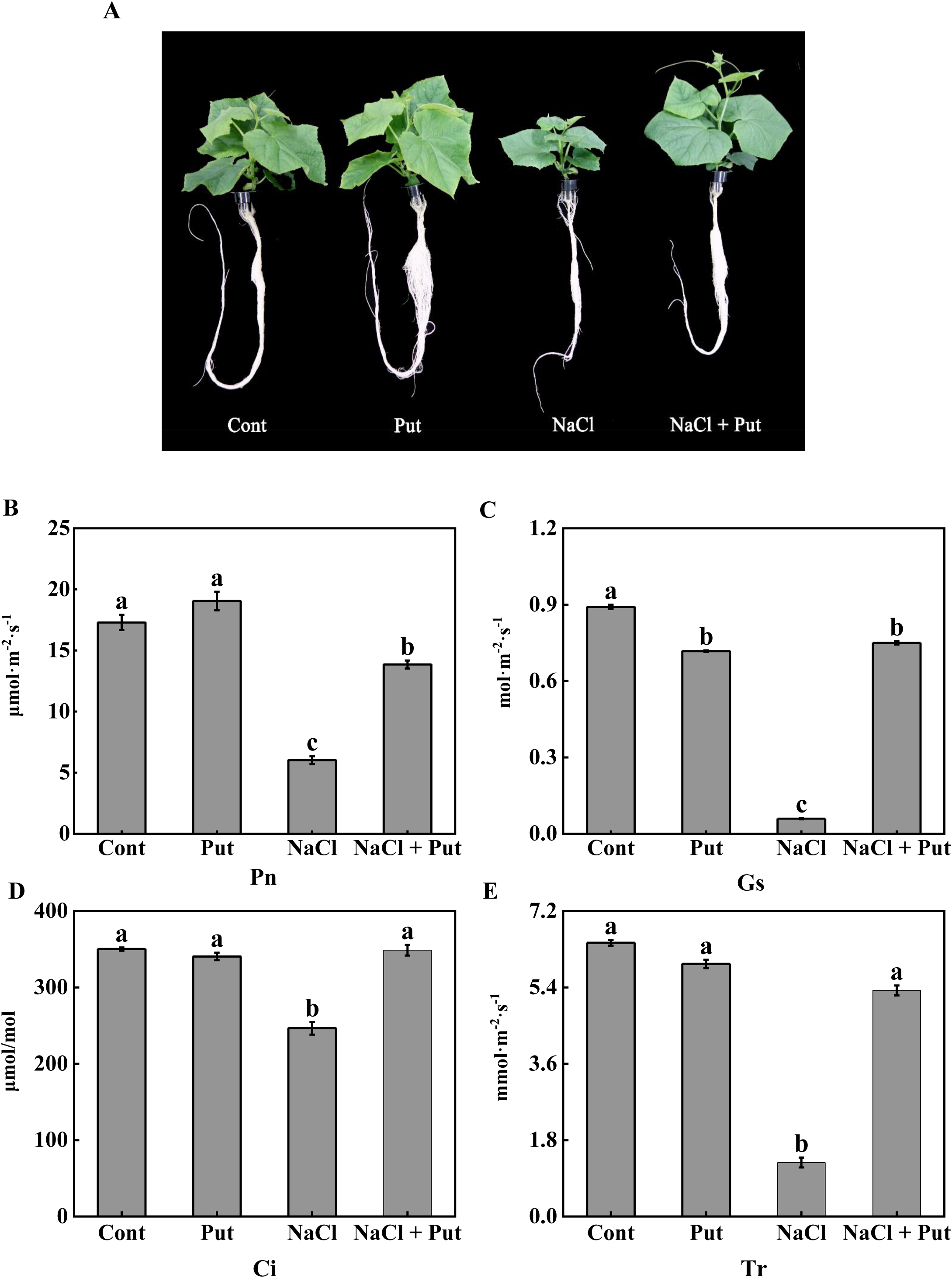
Effects of salt stress and exogenous Put on phenotype and photosynthetic parameters. (A) Growth situation of cucumber seedlings after 7 days treatment with NaCl and/or Put. (B-E) Gasexchange parameters. The third leaves (top to down) in cucumber seedlings after NaCl and/or Put treatment for 7 days were used for chlorophyll measurement. Bars represent the mean ± SD of at least three independent experiments. Different letters indicate significantly different values (P<0.05) by Tukey’s.

### Effects of salt stress and exogenous Put on photosynthetic properties of PSII

To investigate how the PSII performed in salinity and how it affected by Put in growth light of cucumber seedlings, *Fv/Fm*, Y(II), Y(NPQ) and Y(NO) (Y(II)+Y(NPQ)+Y(NO)=1) was detected under 396 μmol m^−2^ s^−1^ light intensity. As shown in Fig. 2A, *Fv/Fm* was similar in all of the treatment plants after 7 days of salt stress, indicating that PSII did not suffer a serious damage under 90 mM NaCl stress condition. Significantly, the effective quantum yield of PSII (Y(II)) in salt stressed cucumber leaves was decreaed by 30% compared with non-stressed cucumber leaves, with the addition of exogenous Put, the Y(II) increased to 1.34 times as high as control. Y(NPQ) is regulatory thermal dissipation quantum yield, which reflects self-protection ability of PSII, Y(NPQ) of NaCl treated cucumbers was 20% higher than Control plants, however it was a little bit lower after spraying exogenous Put under salt stress compared with control. In contrast to Y(NPQ), Y(NO), the non-regulatory thermal dissipation, represents impaired level of PSII, showed no difference in all of the treatment plants, indicating that 90 mM NaCl was not destructively harmful to PSII.

**Fig. 2.**
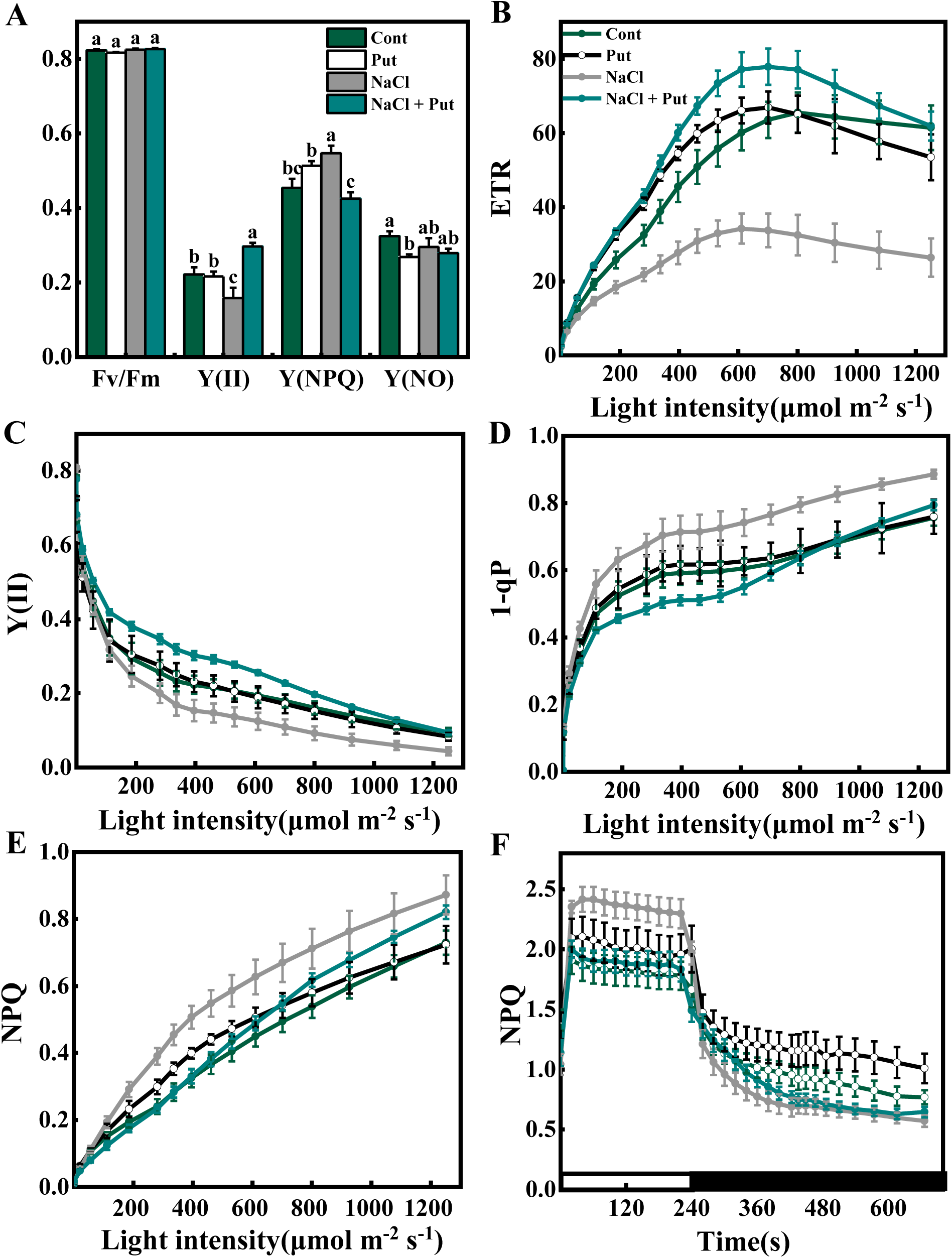
Effects of salt stress and exogenous Put on photosynthetic properties of PSII in cucumber seedlings. The third, fully dark-adapted leaves (top to down) of cucumber seedlings after NaCl and/or Put treatment for 7 days were used in this experiment. (A) *Fv/Fm*, Y(II), Y(NPQ), Y(NO). (B-E) Photoresponse curve of ETR, Y(II), 1-qP and NPQ respectively. (F) Dark-light transition (240 s) and relaxation (506 s) of NPQ. The means ± SD measured at least three independent experiments.

Character of chlorophyll fluorescense are known be responsive to light intensity, so light response curves were measured. Photosynthetic electron flow (electron transport rate, ETR) of NaCl stressed plants was dramatically lower than that in the control plants, but the ETR in NaCl together with Put treated seedlings was even higher than that in the control plants (Fig. 2B). The fast light response curve of Y(II) had a similar trend with ETR curve, it is apparent from this graph that exogenous Put was more obviously beneficial for salt stressed leaves in low light intensity (Fig. 2C). Redox state of Q_A_ (1-qP), the primary electron acceptor of PSII, displayed an obvious increase in salt-treated plants, and it was a little bit lower in NaCl + Put than Cont and Put under moderate light condition (Fig. 2D). NPQ (non-photochemical quenching) is an important photoprotective mechanism to dissipate excess energy, which reflects the self-protected ability, and it could also indirectly represent the utilization ability of light energy. Induction of NPQ in increasing light intensity condition was much stronger in salt-stressed leaves than the control plants, however, in NaCl + Put treatment plants the induction of NPQ was gentle under low light condition, but with the increase of light intensity the induction rate accelerated, and it exceeded Cont and Put treatment plants when light intensity was higher than 400 and 700 μmol m^−2^ s^−1^ respectively (Fig. 2E). These results indicated that 90 mM NaCl stress influenced the photosynthetic properties even in low light condition, and exogenous Put could alleviate the changes caused by salt stress only under low light illumination.

NPQ induction during light to dark transition was also monitired (Fig. 2F). In all treatments, NPQ was transiently induced within 1 min light (396 μmol m^−2^ s^−1^) and relaxed within 4 min dark. NaCl stressed cucumbers exhibited a distinct increase in the rapid induction and relaxation of NPQ, which reached 2.46 after 1 min illumination, and relaxed efficiently, NaCl + Put treatment had a similar dynamic curve with Cont and Put. Photosynthetic systems will drive non-photochemical quenching when they are subjected to salt stress in light, but exogenous Put will decrease the dissipation which is agreement with findings of other groups (Lütz et al., 2005; Ioannidis et al., 2006), that reveals another photoprotection mechanism drived by Put.

### Effects of salt stress and exogenous Put on P700 activity

P700^+^ can absorb 820 nm light when it is oxidated, thus the redox state of P700 was indicated by 820 nm-reflection (Munekage et al., 2004). However, besides P700^+^, other plant tissue can also absorb 820 nm light, and changes of leaves tissue structure will impact measurement of reflection at 820 nm. In order to exclude other influential factors, MR/MR_0_ was used to express reflction kinetic curve at 820 nm light (MR_0_, the first reliable MR at the beginning of illumination) (Strasser et al., 2010).

Figure 3A is a complete curve of 5 times repeat experiment, in which leaves were illuminated on 1 s red light followed with 10 s far-red light. This curve presented a significant decrease in the content of P700 that had potential to be oxidated (the lower MR/MR_0_ signal means the higher P700 activity) in salt-stressed leaves, to a degree, spraying Put was able to protect P700 from salt stress. Moreover, content of oxidative P700 of salt stressed leaves reached to a steady state at the first time illumination of far-red light, but other oxidative curves were all sloped down, which demonstrated that every time after illumination with far-red light, the content of oxidative P700 increased progressively, suggesting that they all stored a number of P700 for urgent condition except treatment NaCl.

**Fig. 3.**
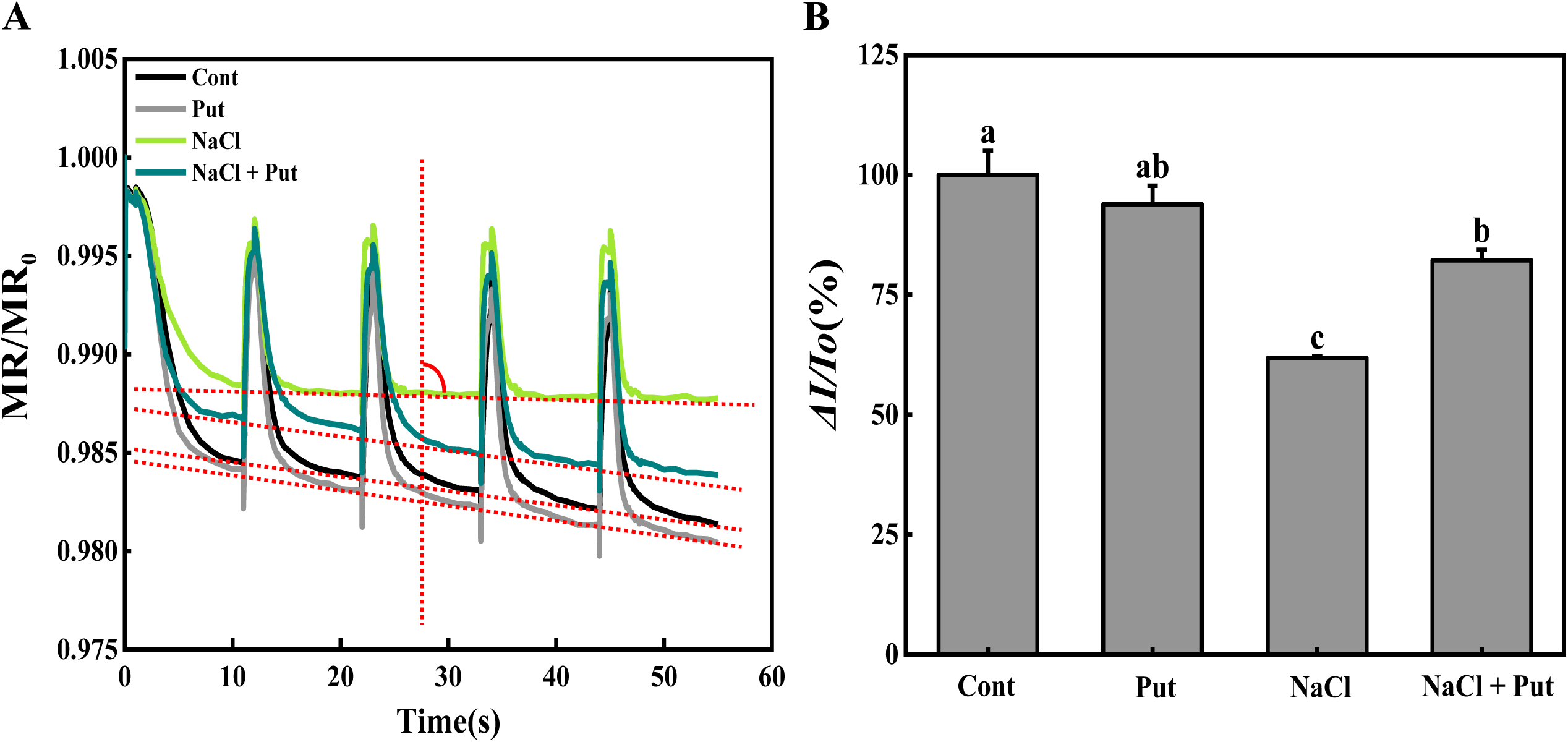
Effects of salt stress and exogenous Put on P700 in cucumber seedlings. (A) 5 repeat measurement of P700 redox state by 1s red light followed with 10 s far-red light illumination. (B) The content of active P700 (*ΔI/Io*). The leaves used for measurement were the same as those used for the Chl fluorescence in Fig. 2.

Relative activity of P700 was also quantified by *ΔI/Io* (P700 relative activity of Cont was assumed to 100%). As shown in Fig. 3B, *ΔI/Io* in salt stressed leaves dropped off by 38.18% than control plants, indicating PSI was seriously inhibited under salt stress. However, *ΔI/Io* in salt stressed with exogenous Put application only decreased by 17.81% in comparison to control plants. Then, methyl viologen (MV), an electron acceptor of P700 down-stream and also an inhibitor of CEF, was used in the experiment, as shown in Fig. S4 when the P700 could be oxidized sufficiently by MV, the content of active P700 in salt stressed cucumber leaves returned to normal (the same as control). Interestingly, exogenous Put obviously increased the P700 activity in non-stressed cucumber.

### Induction of CEF by salt stress and exogenous Put

If indeed P700 is affected by salt stress and exogenous Put application, one would expected to see an effect on the CEF level, therefore CEF was detected by different methods.

Post-illumination is a signal of the transient increase of dark-level chlorophyll fluorescenc after actinic light illumination, which is an important indicator of CEF around PSI (Shikanai et al., 1998). After seedlings were dark-adapted adequately post-illumiation was measured on 396 μmol m^−2^ s^−1^ light intensity. The curves of Fig.4 is fluorescence re-increase condition, it revealed a feeble increase in control cucumber leaves, while exogenous Put could slightly enhance the increase of fluorescence. Significantly, the intensity of fluorescence re-increase signal was dramatically enhanced after NaCl treatment. Interestingly, a much higher CEF level appeared in treatment NaCl + Put plants.

**Fig. 4.**
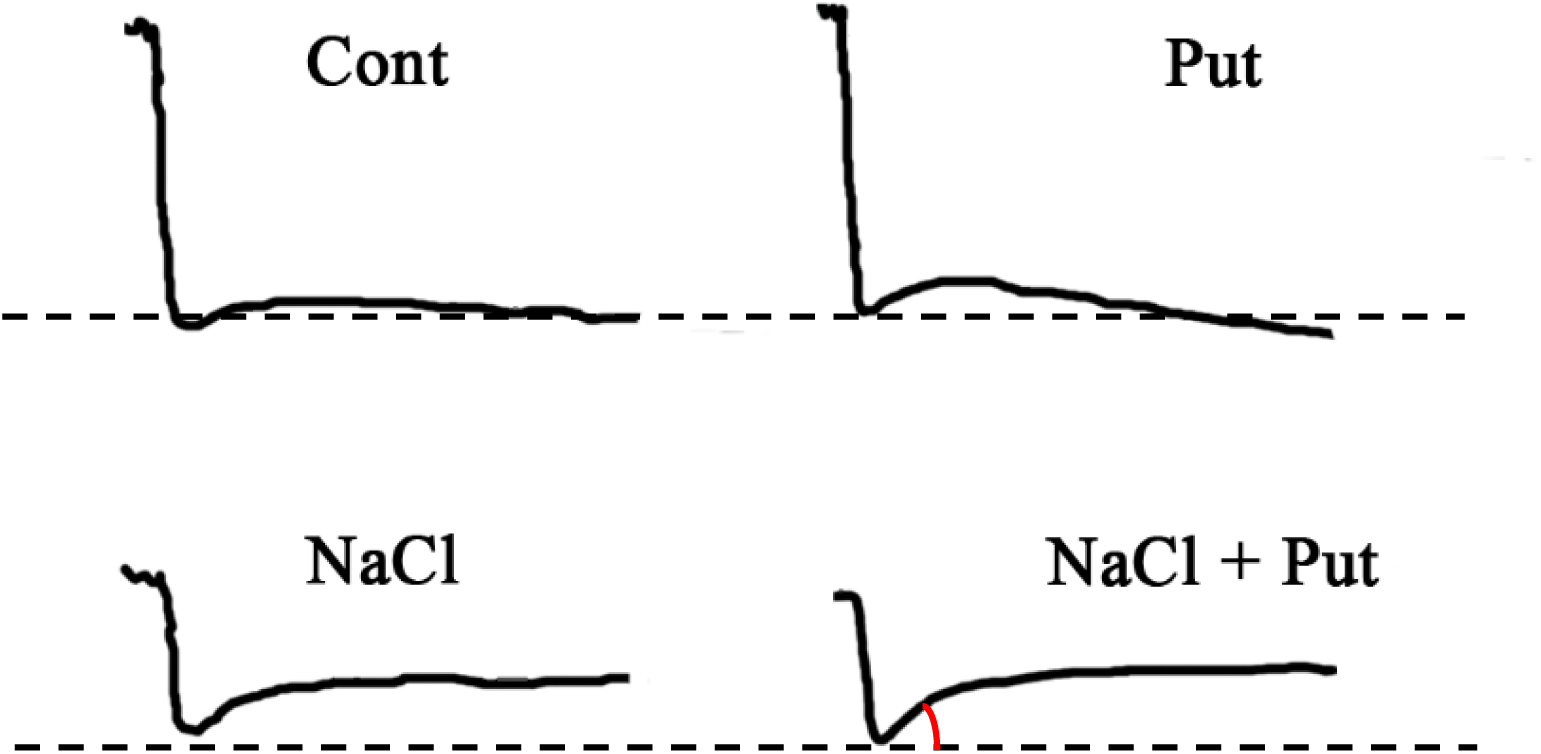
Effects of salt stress and exogenous Put on post-illumination. The typical post-illumination transient increase induction curve was measured by the third (top to down), fully dark-adapted leaves. When the fluorescense got steady, turned off the actinic light and record fluorescense increase signal.

When dark adapted leaves exposed to light, the oxidation rate of their P700 could reflect CEF level, the slower of P700 oxidation rate means the faster of CEF induction rate. So the leaves were illuminated with saturate far-red light for 10 s, meanwhile reflection signal was recorded. Re-reduction of oxidative P700^+^ is another indicator of CEF, a faster re-reduction rate means a faster CEF induction rate. Hence, kinetic curve of P700 oxidation and P700^+^ re-reduction was measured and the half time when the curve got steady was calculated. The re-reduction rate in NaCl and NaCl + Put treatment plants was higher than that in Cont and Put treatment plants (Fig. 5A), the *t_1/2_* of P700^+^ re-reduction was 0.21, 0.20, 0.16, 0.11s in Cont, Put, NaCl, NaCl + Put respectively (Fig. 5B). In addition, the oxidation rate of P700 had a stepwise increase from Cont to NaCl + Put (Fig. 4C). The *t_1/2_* of P700 oxidation was 2.18, 2.20, 2.34, 2.61s in Cont, Put, NaCl, NaCl + Put respectively (Fig. 4D). These results suggested that salt stress induced CEF to protect the photosynthetic apparatus, and exogenous Put acted as an accelerator to enhance CEF induction.

**Fig. 5.**
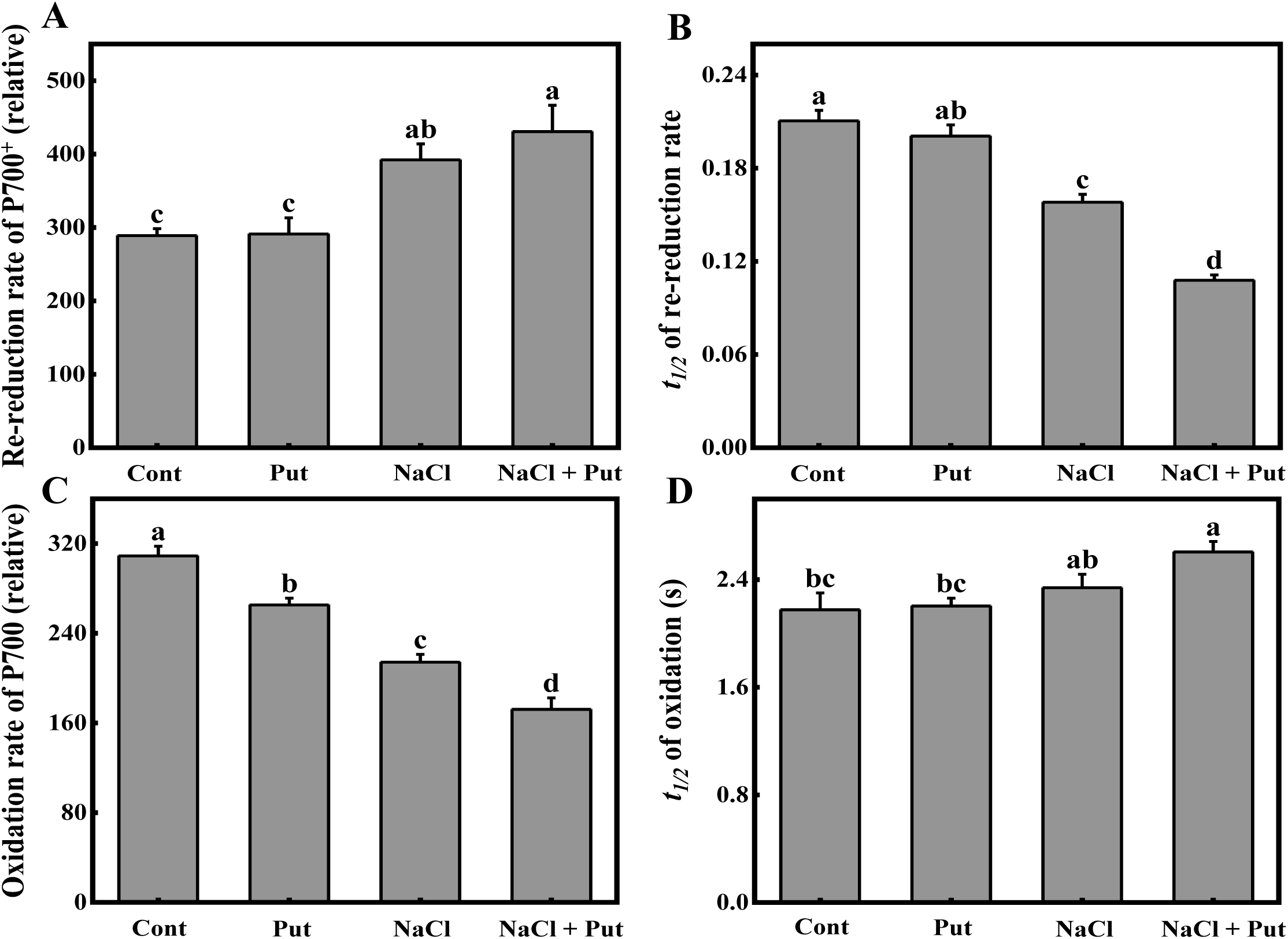
The redox rate and *t_1/2_* of P700 in different treated cucumber leaves. Bars represent the mean ± SD of at least three independent experiments. Different letters indicate significantly different values (P<0.05) by Tukey’s test.

### NaCl stress and Put treatment changes pmf formation across thylakoid membrane in cucumber leaves

Due to changes of electron transport chain could influence H^+^ accumulation in lumen, we measured *trans*-thylakoid proton gradient (ΔpH) by delayed fluorescence (Fig. 6A). Increase from D_2_ to I_4_ accompanied by excitaiton of PSI after illumination, the amplitude is connected with formation of ΔpH (Evans and Crofts, 1973). It should be noted that the detector can obtain various induction curve when under different light intensity condition. 3000 μmol m^−2^ s^−1^ light induced an inconspicuous I_4_, however, the I_4_ induced by 600 μmol m^−2^ s^−1^ light was apparent. (I_4_-D_2_)/D_2_ is a representation for ΔpH, salt stressed cucumber leaves induced high level of CEF with high ΔpH. Although CEF intensity of NaCl + Put-treated plants was higher than that in NaCl only treated plants, the ΔpH was decreased with Put application under salt stress (Fig. 6C). With our results, the changes in one more characteristic point, I_1_, was observed. This first maximum is related to the accumulation of *trans*-membrane electrical potential (Δψ) by oxidation of PSI (Pospíšil and Dau, 2002). The maximum I_1_ in salt stressed leaves was lower than other treatments (Fig. 6A), indicating a lower gradient of Δψ in salt stressed leaves.

**Fig. 6.**
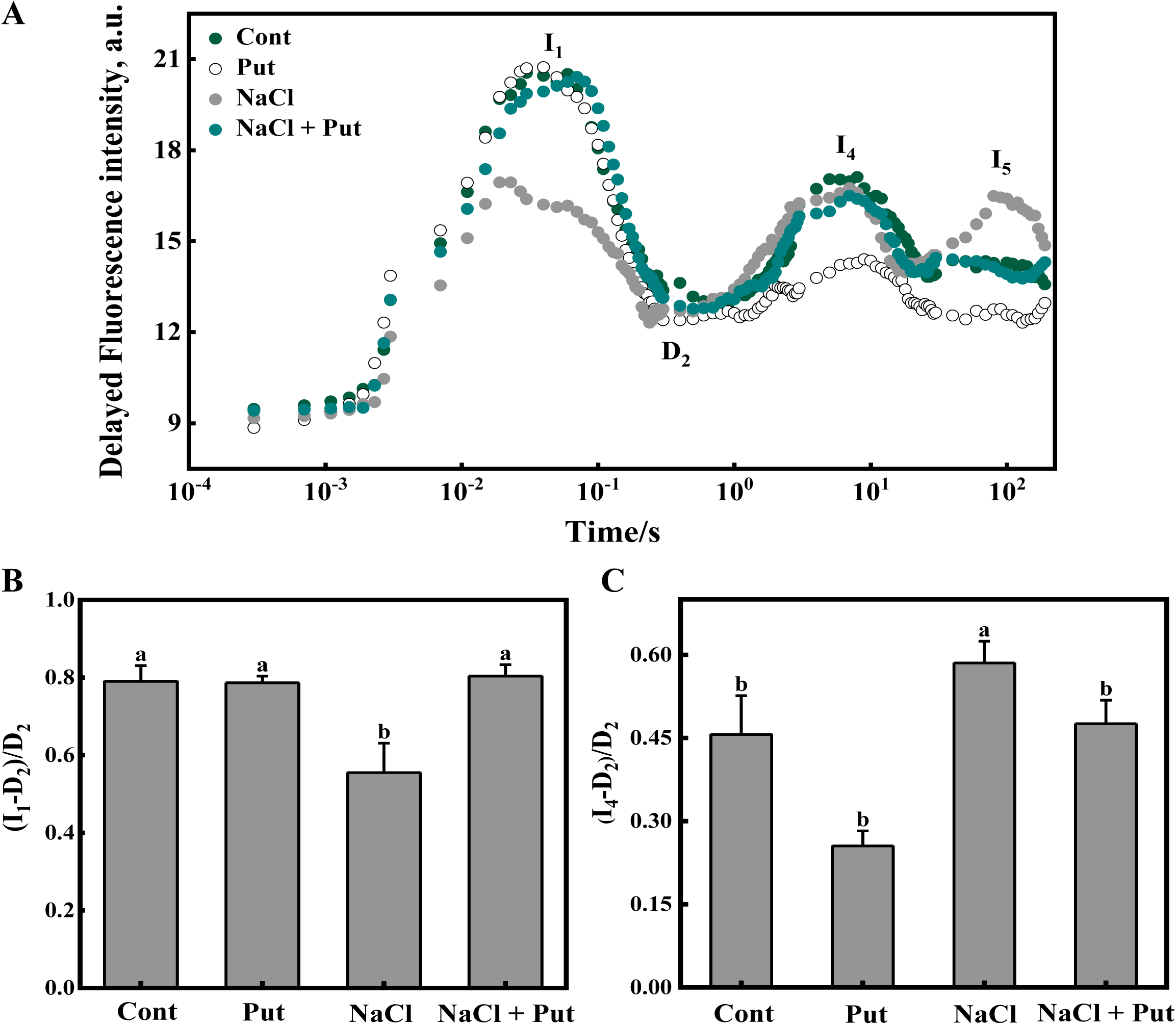
Effects of salt stress and exogenous Put on delayed fluorescence. (A) Delayed fluorescence induction curves (on log time scale), maxima in the figures are designated as I_1_, I_4_, I_5_, while the minimum is designated by D_2_. Other maxima are not pronounced in our samples. (B, C) Delayed fluorescence parameters. Bars represent the mean ± SD of at least three independent experiments. Different letters indicate significantly different values (P<0.05) by Tukey’s test.

### Effects of salt stress and exogenous Put on light induced ATP, ATP/NADPH, NADP^+^/NADPH and RCA activity in cucumber leaves under different treatments

ΔpH together with Δψ comprises proton motive force (*pmf*) which fuels ATP generation, ATP and NADPH are production of photosynthetic electron transport, act as assimilatory power for CO_2_ fixation. Therefore, content of ATP, NADPH and NADP^+^ is another indirect important indicator for electron transport activity. They are all measured in light- and dark-adapted leaves, light induced production was calculated as content in light minus in dark. Fig. 7A revealed a significant decrease of ATP production induced by light in salt-stressed leaves, expectedly, exogonous Put promoted the ATP generation 40.36% more than that of treatment NaCl. In non-stress conditions, the ATP/NADPH ratio was maintained to 0.33 for plants normal growth, however, this ratio decreased to 0.09 after 7 days of salt stress (Fig. 7B). The reason why the ATP/NADPH ratio decreased such violently not just because of decline of ATP production, but also due to the depressed RCA (rubisco activase) activity (Fig. 7D) making the NADPH unable to be consumed efficiently in carbon assimilation phase, consequently resulting in accumulation of NADPH, further leading to an extremely low NADP^+^/NADPH ratio (Fig. 7C), these all manifest that the thylakoid membrane was over-reduced in salt-stressed cucumber plants.

**Fig. 7.**
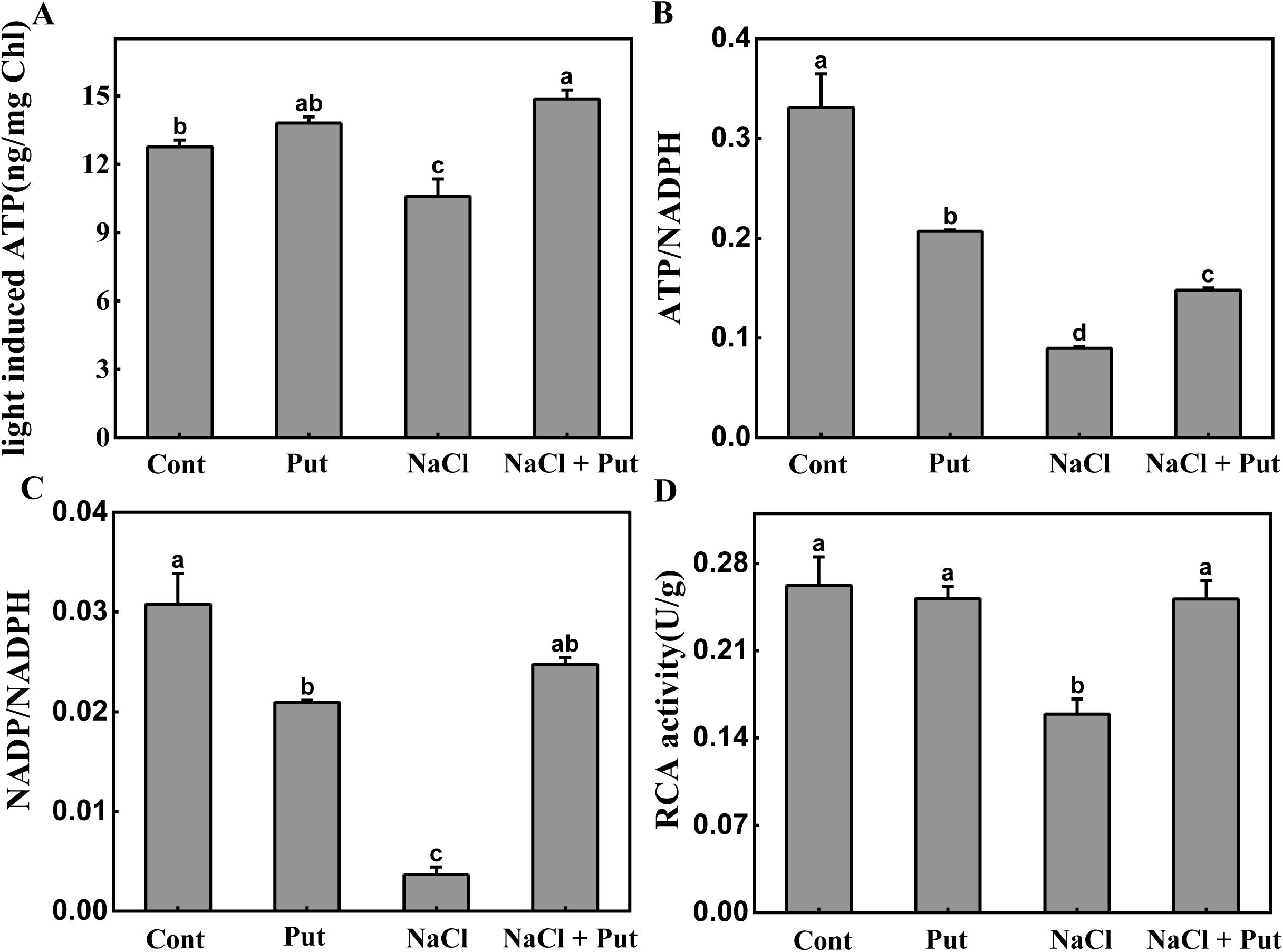
Light induced ATP, ATP/NADPH, NADP/NADPH and RCA activity in different treated plants. The dark-adapted and light-induced leaves were used for these measurements. (A) Production of Light induced ATP; (B) ATP/NADPH; (C) NADP/NADPH; (D) Activity of RCA. Different letters indicate significantly different values (P<0.05) by Tukey’s test.

### Changes of gene expression analysis of electron transport related proteins

It has been demonstrated that, ferredoxin 2 in Arabidopsis is a dominated protein of electron transport chain, which participates in LEF, and Ferredoxin 1 is an accessory protein attributed to CEF (Holtgrefe et al., 2003; Blanco et al., 2011; Liu et al., 2013). We compared the amino acid sequence in cucumber with that in Arabidopsis. The sequence of Fd L-A like protein in *Cucumis sativus* has a high homology with Fd1 in arabidopsis, the similarity reaches to 64%, and the amino acid sequence of Fd in Cucumis sativus is 63% similar to that of Fd2 in arabidopsis (Fig. S3). As expected, in salt stress condition the transcript level of Fd in cucumber leaves was significantly down-regulated (30.27% lower than Cont), and the level of Fd L-A like was up-regulated (136.68% higher than Cont), however, exogenous Put increased the transcript level of Fd in salt-stressed by 2.47 times compared wtih treatment NaCl, but not affacted transcription of Fd L-A like (Fig. 8B).

**Fig. 8.**
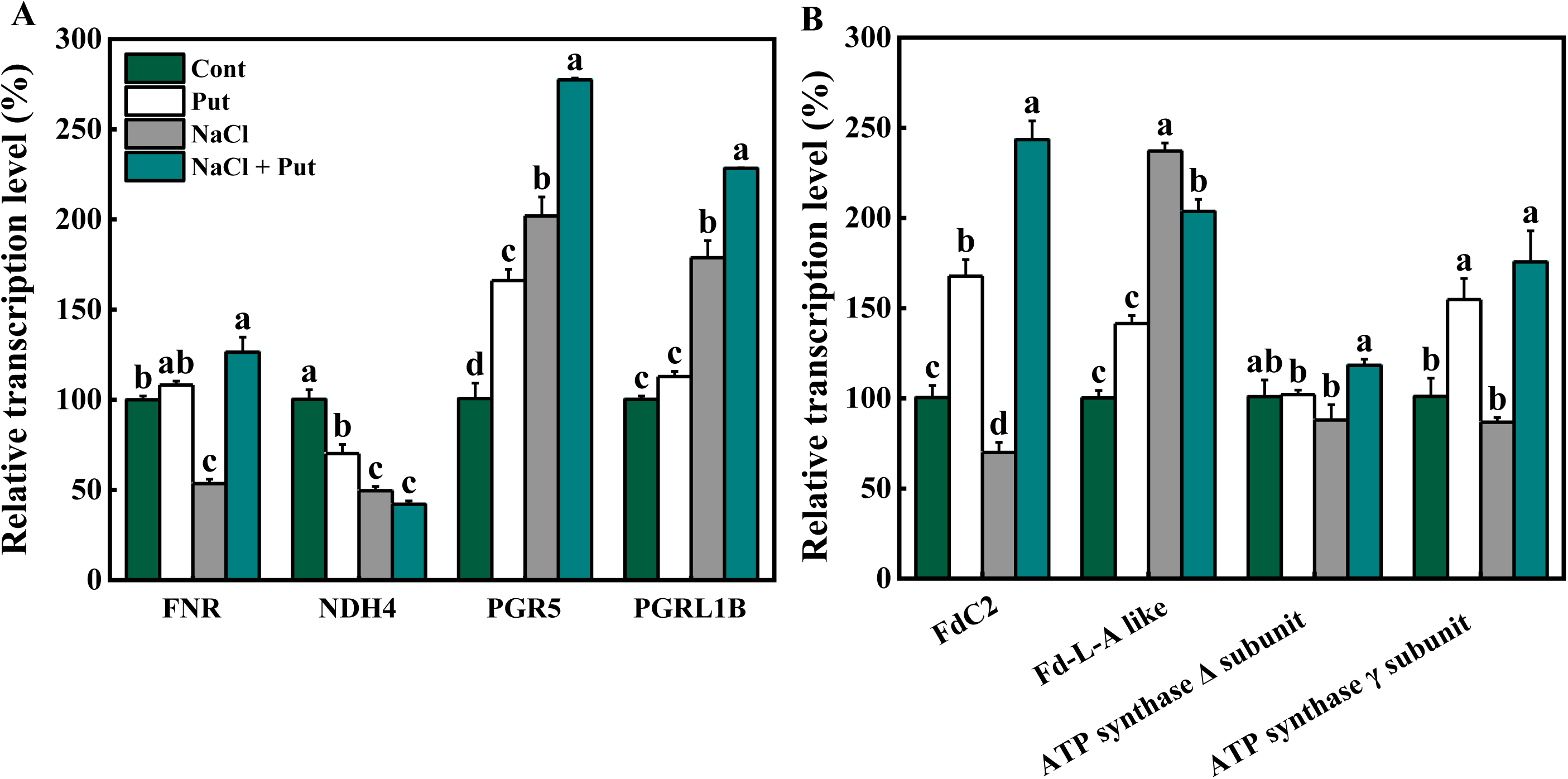
Genes expression analysis of electron transport related proteins in different treated plants. Transcript levels of these genes were measured after NaCl and/or Put treatment for 7 days. Different letters indicate significantly different values (P<0.05) by Tukey’s test.

Fd-NADPH oxidoreductase (FNR) is an electron receptor of PSI in electron transport chain. Some report suggested that FNR was involved in NDH-dependent CEF, some said FNR had no connection with CEF (Zhang et al., 2001; Medina and Gómez-Moreno, 2004). Our gene expression results showed a down-regulation of FNR in response to salt stress in cucumber leaves, but the up-regulation was pronounced after spraying with Put in salt-stressed cucumber leaves (Fig. 8A).

The gene expression level of the dominating proteins regulationg the two known CEF pathways were also determined. NDH4 is a main subunit of NDH complex in cucumber chloroplast, unexpectedly its transcript level was down-regulated in salt stressed leaves, and which performed similar to treatment NaCl + Put. In contrast to NDH4, transcript levels of PGR5 and PGRL1 were distinctly up-regulated in response to salt stress (100.42% and 78.24% for PGR5 and PGRL1 respectively higher than Cont), even more, exogenous Put induced higher transcription level of PGR5/PGRL1 than NaCl treatment. This results proved that the PGR5/PGRL1-dependent pathway is the major CEF in response to salt stress (Fig. 8A). Accordingly, transcript levels of ATP synthase Δ and γ subunit were all down-regulated after 7 days of salt stress, and it was unchanged even up-regulated in treatment NaCl + Put compared with Cont (Fig. 8B).

## Discussion

### NaCl stress induced a serious photoinhibition rather than a photodamge in PSI and PSII of cucumber leaves

Effects of salinity on two photosystems were controversial all the time, due to different plant species and various treatment protocol. Some studies proved that PSII was sensitive to salinity (Demetriou et al., 2007; Mehta et al., 2010), whereas other studies demonstrated a high salt-resistance of PSII (Lu et al., 2003; Yan et al., 2015). In this study, 90 mM NaCl was no effect on *Fv/Fm* (Fig. 2A), which was in accordance with previously study (Chen et al., 2013), and the Y(II) of salt stressed cucumber leaves relaxed efficiently after light turned off, although it was significantly lower than other treatments in the light-induce phase (Fig. S2). Furthermore, the Y(NO) didn’t change after salt stress. These results suggested that 90mM NaCl stress would not cause an irreversible damage on PSII, which was consistent with the changeless of chlorophyll content (Fig. S1D), and the excess excited energy that can not be utilized was consumed by non-photochemical quenching (Fig. 2A, E, F) to protect PSII from photodamge.

In contrast to PSII, few studies focused on how salt stress influences PSI capacity in plants. The electron flow from PSII is essential for PSI photoinhibition, if electrons are blocked before PSI, PSI photoinhibition can be supressed and it is helpful for PSI recovery (Zhang et al., 2011). Unfortunately, PSII was not efficiently inhibited in our study, excess electrons rushed into PSI, resulting to over-reducing in the accept side of PSI (Fig. 7C) and the active of P700 was seriously impaired (Fig. 3). Nevertheless, the salt-influence of P700 was not irreversible, when MV accepted electrons from P700, the oxidizable P700 activity was recovered (Fig. S4).

Taken together, these results indicated that 90 mM NaCl was not acutely harmful to PSII and PSI, it would inhibit the activity of the two photosystems, but not damage them.

### Exogenous Put induces a stronger CEF under salt stress

The results showed that NaCl stress compelled the leaves to drive a high level of CEF (Fig. 4, 5), which was effective to protect PSII, but not enough for PSI. Increasing CEF induction by exogenous Put was apparent in our results. PAs were seen by researchers as cations and chemical equilibrium buffers (Ioannidis et al., 2006). Cationic effects can induce stacking of thylakoids that similarly to divalent inorganic cations. Buffering role of PAs can stimulate ATP synthesis (Ioannidis and Kotzabasis, 2007). Put drastically stimulated phosphorylation in light, the ATP generated in NaCl + Put treatment was 40.40% more than that in NaCl single treated leaves. In addition, PAs also participate in the modulation of *pmf* in thylakoid *in vivo* by dissipating ΔpH and favoring Δψ (Ioannidis et al., 2012; Ioannidis and Kotzabasis, 2014). A lower ΔpH in Put treated plants was shown in Fig. 6C. Abundant ATP is essential for PSI recovery, that means more active P700 can participate in CEF (Fig. 3) without considering over-acidification in lumen. Therefore, CEF induction mechanism in treatment NaCl + Put basically benefits from buffering character of Put.

### Enhanced CEF promotes ATP production

In return, increased level of CEF supplied more ATP for CO_2_ assimilation in NaCl + Put (Fig. 7A). Inhibition of CO_2_ assimilation by stresses leads to over-reduction of electron transport chain indirectly, increasing reduction level of intersystem electron transporter promotes CEF around PSI (Suorsa et al., 2015). Indeed, salt stressed leaves had a higher redox state of PQ pool (represented by 1-qP, Fig. 2D) and was over-reduced in PSI acceptor side (Fig. 7C), which was able to be an important factor to induce a higher CEF in NaCl stressed leaves (Fig. 4, 5). Additionally, ATP content is also a regulatory factor for interconversion between LEF and CEF. Salt stressed leaves with a lower light-induced ATP content (Fig. 7A) needs an alternative pathway such as CEF around PSI to supplement the deficiency, which used to maintain themselves alive and continue to growth. Whereas, exogenous Put played a critical role like a fuel, drived a stronger and completely different salt tolerance mechanism in thylakoid.

Sufficient ATP together with surplus NADPH contributes to CO_2_ assimilation progress, the activity of RCA detected in NaCl + Put treatment was higher than that in NaCl treated leaves (Fig. 7D), which directly helpful to a rebalance of NADP^+^/NADPH ratio (Fig. 7C). Increased number of NADP^+^ relieves pressure at PSI acceptor side, and a higher level of CEF makes excess electrons revolved around PSI also alleviate stress at PSI accepter side.

### Energy dissipation is switched from non-photochemical quenching to photochemical quenching after Put treatment for salt stressed plants

If the flow of electrons through the electron transport chain exceeds the capacity of metabolism to consume the reductant production, then potentially harmful side reactions are liable to occur (Hald et al., 2008). The best characterised regulatory mechanism for limiting damage is non photochemical quenching (NPQ) (Li et al., 2000; 2002; Pascal et al., 2005). According to relaxation kinetics in darkness following a period of illumination, the energy-dependent non-photochemical quenching (qE) is the major and most rapid component of NPQ in plants (Fig. 2F; (Horton and Hague, 1988). Generation of qE requires the build-up of trans-membrane ΔpH, which in the chloroplast is mainly induced by electron transport chain (including LEF and CEF) (Aihara et al., 2016; Sun et al., 2017). ΔpH together with Δψ composes *pmf* in thylakoid lumen, and they can be converted to each other (Avenson et al., 2004; 2005). When plants are under optimal condition, the down-regulation of photochemical quenching is not needed, so a large fraction of *pmf* can be stored as Δψ, leading to moderate lumen pH and low qE, even at high *pmf* (and thus high rates of ATP synthesis). When plants suffer from environmental stresses e.g., salt stress, high temperature, high light, chilling et al., the photoprotection is obligatory, *pmf* can be predominantly stored as ΔpH, maximizing lumen acidification for a given *pmf* (Ioannidis et al., 2012). In the present study, we found that stressed leaves showed lower Δψ and higher ΔpH compared with the control plants (Fig. 6 A and C), which allowed the membrane to induce strong qE and down-regulate photochemical quenching, resulting in the production of ATP slowed down. Nevertheless, over-acidification in lumen will inhibit the oxidation of plastoquinol by deactivating the Cyt *b_6_f* (Harbinson and Hedley, 1993; Laisk et al., 2005). Exogenous Put neutralized excess H^+^ in lumen and built a moderate pH condition, leading to maintain high level of CEF in salt stressed leaves, no need to worry about inactivation of electron transporters. Furthermore, the qE was reduced, and most of the excited energy were transferred to photochemical quenching. The efficient photochemical electron transport chain (PETC) in NaCl + Put treated plants is necessary for normal growth (Fig. 1 and S1); High level of CEF is driven to produce extra ATP for PSI repair and CO_2_ fixation, and it plays a buffer role in PSI accepter side (Fig. 9).

**Fig. 9.**
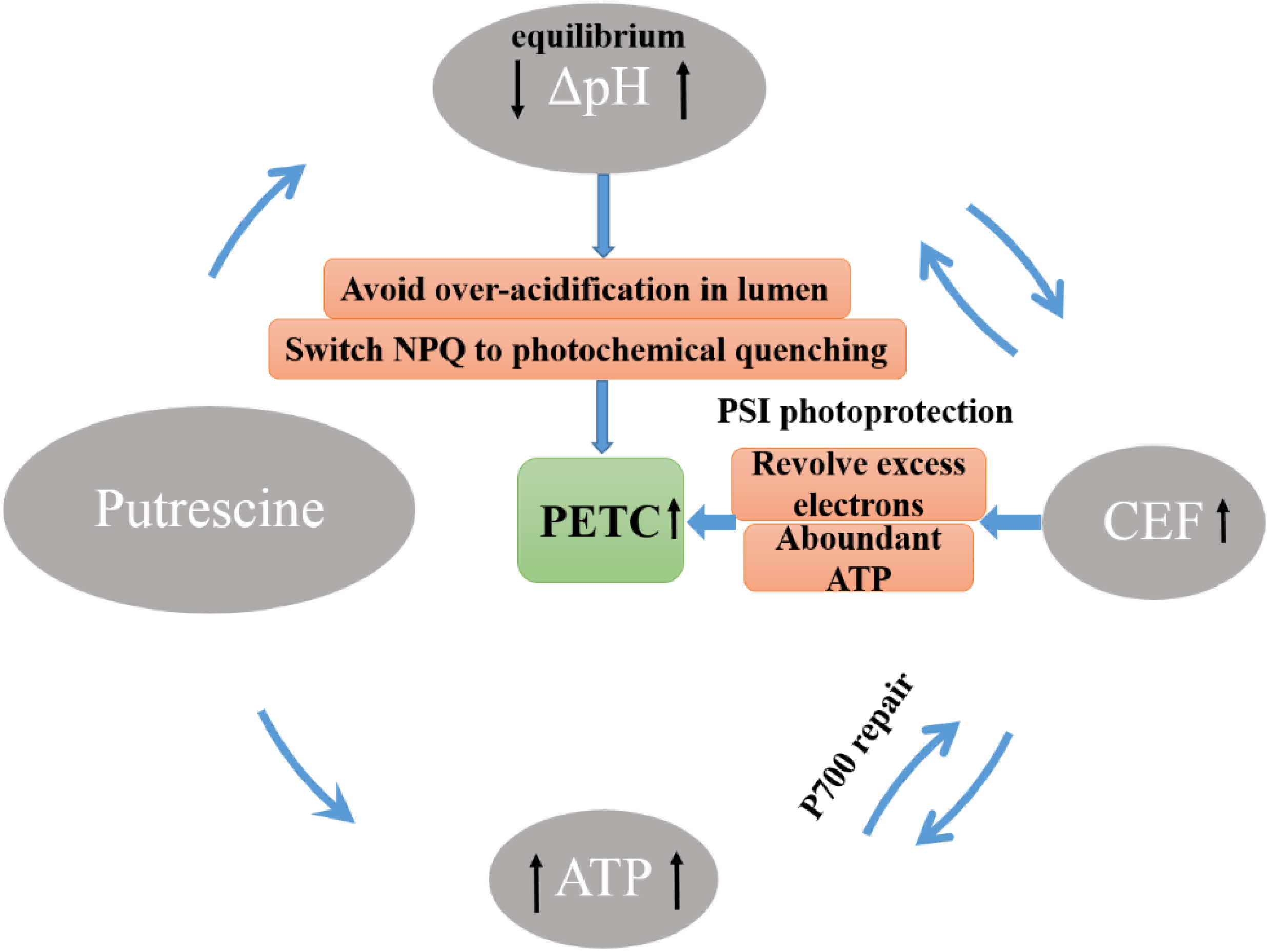
Model for putrescine inducing photoprotection in cucumber leaves under salt stress. Exogenous putrescine reduces ΔpH across thylakoid membrane to avoid over-acidification in lumen, accordingly decreased nonphotochemical quenching. In addition, ATP synthesis is accelerated by Put. The lower ΔpH and higher content of ATP are crucial for strengtherning CEF to enhance photoprotection.

In conclusion, 90 mM NaCl caused photoinhibition of both photosystems, CEF is induced to protect them not be damaged. Exogenous Put reduces ΔpH across thylakoid membrane to avoid over-acidification in lumen, accompanied by decreasing non-photochemical quenching. In addition, ATP synthesis is accelerated by Put. The lower ΔpH and higher content of ATP are crucial for strengthening CEF to enhance photoprotection of thylakoid apparatus, further increasing the utilizaiton efficiency of light energy (Fig. 9).

## Supplementary data

Supplementary data are available at *JXB* online.

Table. S1. Primer sequences used in quantitative real-time PCR assays

Figure. S1. Changes of growth parameters in cucumber seedlings under NaCl and/or Put treatment for 7 days.

Figure. S2. Dark-relaxation of Y(II) of cucumber leaves under NaCl and/or Put treatment for 7 days.

Figure. S3. Sequence alignment of ferredoxin related proteins in *cucumis sativus* L. and A. *thaliana.*

Figure. S4. The redox state of P700 in cucumber seedlings when the leaves there treated with 2 mM MV (melthyl viologen, an electron acceptor of P700, also an inhibitor of CEF).

## Acknowledgements

Thanks to my supervisor, Guo Shirong, for his significant support for my work. And I am grateful to the numerous individuals who participated in this research. Mr. Sheng Shu and Mr. Yu Wang provided critical discussion and comments, I also thank Ruonan Yuan for help with experiment technology. This work was supported by the National Natural Science Foundation of China (No. 31672199 and No. 31471869) and was supported by China Agriculture Research System (CARS-23-B12).

